# Co-occurrence, habitat use and activity pattern of carnivore species in a coastal area of Argentina

**DOI:** 10.1101/2020.03.04.976415

**Authors:** N.C. Caruso, E.M. Luengos Vidal, M.C. Manfredi, M.S. Araujo, M. Lucherini, E.B. Casanave

## Abstract

Land-sea interface is an ecotone where the intersection of marine and terrestrial ecosystems create unique ecological conditions for terrestrial mobile species and freshwater-adapted organisms to exploit marine-derived food resources. Mammalian carnivores play an important role in almost any ecosystem where they live due to their top-down (or trophic cascade) effects on prey species and primary producers, thus structuring ecosystems along varied food-web pathways. We use camera trapping to study the patterns of coexistence, habitat use and activity pattern of carnivores species in a coastal area in southern Buenos Aires province, Argentina. We were able to detect five of the seven species of Mammalian carnivores being the Pampas fox *Lycalopex gymnocercus* and Geoffroy’s cat *Leopardus geoffroyi* the two most common. Geoffroy’s cat seems to use more intensively those areas close to the shoreline, while we found little support of it for Pampas fox; which seems to use more inland areas. Congruently, we found evidence of a lack of spatial and, to a lower extent, temporal avoidance between the two most common carnivore species of our study area. Our findings support those previous studies indicating that the coastal dunes have an important role in the conservation of the biodiversity of Buenos Aires province. Wildlife conservation is compatible with carefully-designed ecotourism and limited infrastructure development and this may be a unique chance for the areas of Buenos Aires coast that have not been affected yet by poorly planned, conservation-unfriendly urbanization.

## 1. Introduction

Land-sea interface is an ecotone where the intersection of marine and terrestrial ecosystems create unique ecological conditions. At this ecotone, terrestrial mobile species and freshwater-adapted organisms are able to get into near-shore marine and estuarine waters or inter-tidal and marginal habitats (e.g., supralittoral zone) to exploit marine-derived food resources (Carlton & Hodder, 2003; Lillywhite, Sheehy III, & Zaidan III, 2008). In coastal and island systems, allochthonous marine resources can greatly subsidize terrestrial food webs, especially in areas of low productivity. Indeed, marine inputs represent a resource subsidy that may contributes to higher population densities of terrestrial consumers than those supported by *in situ* terrestrial resources alone (Behrendorff, Leung, & Allen, 2019; Brown et al., 2015; Rose & Polis, 1998).

Mammalian carnivores play an important role in almost any ecosystem where they live due to their top-down (or trophic cascade) effects on prey species and primary producers (Glen & Dickman, 2005; W. J. Ripple et al., 2014; Terborgh & Estes, 2013). Moreover, large carnivores can potentially limit mesocarnivores species through intraguild competition, thus structuring ecosystems along varied food-web pathways (William J Ripple, Wirsing, Wilmers, & Letnic, 2013; Suraci, Clinchy, Dill, Roberts, & Zanette, 2016). Although a wide range of factors can affect carnivore populations, interspecific competition and resource partitioning have been proposed as one of the most important factors structuring carnivore communities (Di Bitetti, De Angelo, Di Blanco, & Paviolo, 2010; May et al., 2008). Thus, the presence of a specie in a given environment is the results of a trade-off between the quality of the habitat and the interspecific competence that directly or indirectly limits the access to resources (Araujo & Guisan, 2006; Soberón, 2007). Within this framework, MacArthur and Levins (1967) proposed that two competitive species can co exists only if their niche differ, at least partially, along one or more dimensions. Later, Schoener (1974) argued that niche segregation between two competitive species is multidimensional and that the spatial (habitat), the trophic, and the temporal are the most common niche dimensions on which two species segregate.

The aim of this study was to gain ecological information about the carnivore community in a coastal area that represents a relict of semi-natural grasslands in southern Buenos Aires province, Argentina. Because the natural grasslands that previously covered almost the entire territory of the province have almost disappeared due to their transformation into croplands, these relicts have a high conservation value (Bilenca & Miñarro, 2004). Thus, our primary objective was to understand the relevance of these coastal habitats for the conservation of the native community of carnivores. Additionally, we aimed to explore if the local carnivore community was randomly organized in terms of species’ coexistence and interactions and if not, what are the main interspecific differences in habitat use and activity patterns. We hypothesized that carnivore community would be comparatively diverse and that the species composing it would show relatively ample niche overlap between each other in terms of habitat and activity pattern due to the greater availability of resources found in this coastal area in relation to the habitats largely modified by anthropogenic activities typical of more inland areas.

## 2. Methods

### 2.1. Study area

Fieldwork was carried out in the “Arroyo Los Gauchos Nature Reserve”, located in the southwestern most part of Buenos Aires province (Coronel Dorrego county, 38°56’1.93”S, 60°45’9.37”O; Figure 1). The “Arroyo Los Gauchos Nature Reserve” is a coastal protected area of 7.07 km^2^ forming part of the Pampas ecoregion. Its habitat is characterized by the presence of dunes, both non-vegetated and covered by psammophyte vegetation, and a general low topography with interdune depressions occupied by small wetland. The climate is temperate, the annual mean temperature is 14.1°C, with a maximum in January and a minimum in July, and the annual mean precipitation is about 850 mm (Celsi & Giussani, 2019; Monserrat, Celsi, & Fontana, 2012).

**Figure 1.**
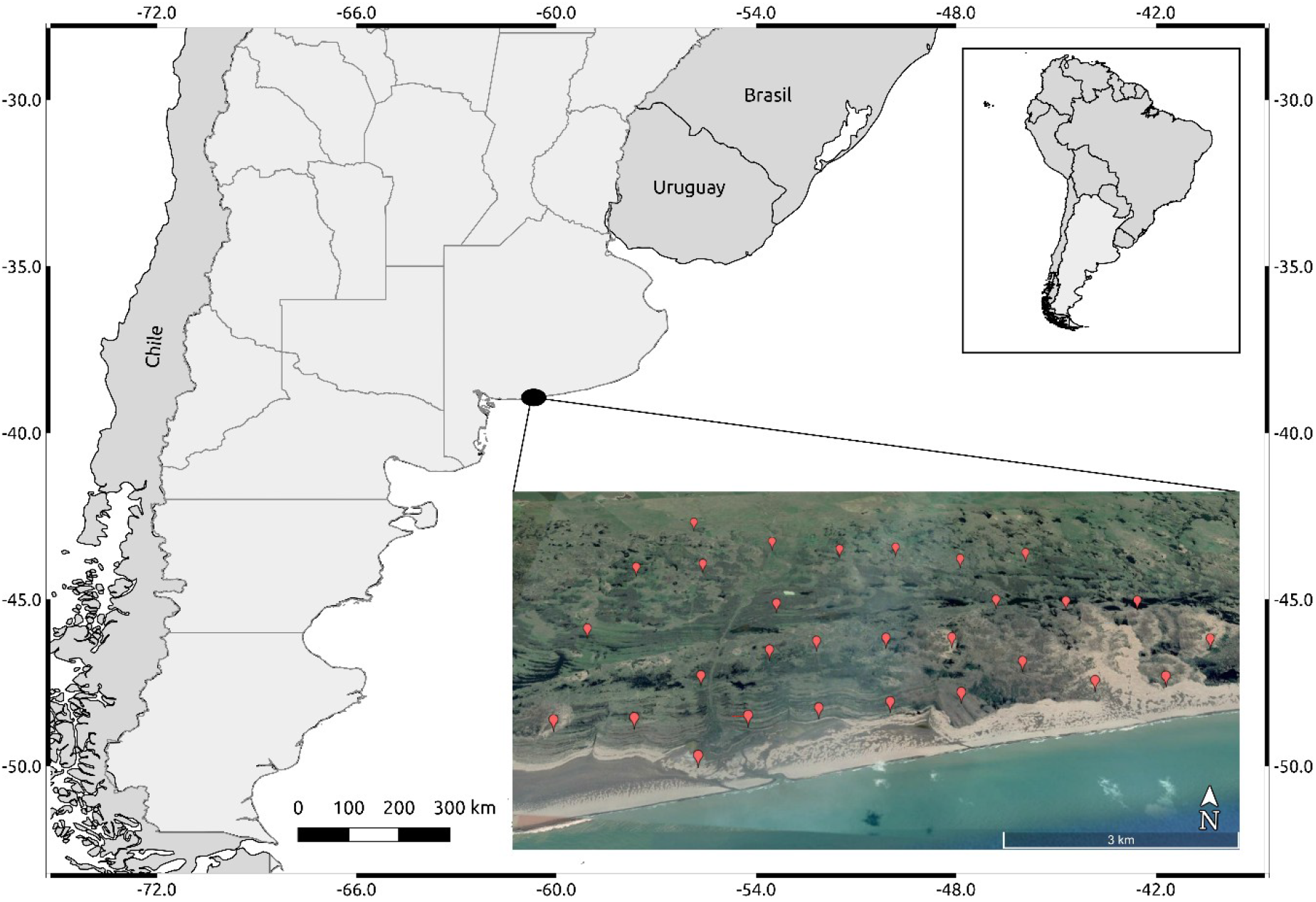
Map of the study area. Black dot in the bigger map shows the relative location of the study area in Argentina. Red symbols in the inner image represent the camera trap sites.

Although the study area is formally protected by the provincial government since 2011, it lacks implementation and its habitats are preserved primarily because of the difficulty of access. Until now, human disturbance has had a low impact in comparison to the other coastal areas of the province. Activities such as cattle farming, forestry (principally *Eucaliptus ssp*. and *Pinus ssp*. plantations) and tourism -which are widespread elsewhere-are still limited and urban development is low (Monserrat & Celsi, 2009; Monserrat & Codignotto, 2013). The area was proposed as Valuable Grassland Area (Bilenca & Miñarro, 2004) well before its establishment as protected area due to its high biodiversity (more than 72 vascular plant species) and the presence of endemic rare or endangered vertebrate species, including a rodent, *Ctenomys australis* and a lizard, *Liolaemus multimaculatus* (Monserrat & Celsi, 2009).

### 2.2. Data collection

We conducted camera trapping from January 2009 to March 2009 using a combination of both film and digital camera traps. The photo-trapping stations (N=29) were located in places where we had previously registered indirect evidence of carnivore activity or along trails. Because the Geoffroy’s cat is considered rare or nearly extinct in most of the anthropogenically modified habitats of the territory of the province (Castillo, Luengos Vidal, Lucherini, & Casanave, 2008), we focused our efforts on this felid. Thus, to reduce the chances of missing the presence of undetected Geoffroy’s cat individuals, the stations were separated by an approximate distance of 1 km. This distance was selected based on the information on home range size and movements of Geoffroy’s cats that were radiotracked in a previous study in the Pampas grassland (Manfredi, Soler, Lucherini, & Casanave, 2006).

Odorous baits (Bobcat Urine and Bobcat Gland Lure) were used at all stations in order to increase detection probability of carnivores, which are frequently elusive species. At each station we deployed one camera that was running continuously and set up to work with the minimal delay between pictures available for each model. All cameras were operational 24 hours per day except in instances of malfunction or damage caused by cattle, climate or other factors. We checked cameras every 5 days to replace batteries, film or memory card and to ensure their proper functioning and we replaced the cameras when they failed to minimize the amount of missing data. Sampling effort was calculated as the sum of the effective sampling days (omitting those days in which the cameras did not work) along the total number of stations (Di Bitetti, Paviolo, & De Angelo, 2006).

The habitat surrounding camera stations was in situ evaluated within a radius of 20 m to visually estimate the proportion of each habitat type. Seven habitat types were identified as the most characteristics in the study area: “BG”, bare ground, mostly non-vegetated dunes or any other portion of land without vegetation cover; “LG”, low grassland, grasses (Poaceae) less than 30 cm tall; “TG”, medium-tall grassland, areas covered by grassy vegetation (Poaceae) taller than 30 cm; “J”, areas covered by *Juncus spp*.; “C”, areas covered by *Cortaderia spp*.; “H”, areas covered by *Hyalis argentea*; “S”, shrubland, mostly dominated by *Lycium chilensis, Baccharis divaricata and Discaria americana*. Additional we measured “D”, the linear distance (m) from each camera station to the shoreline. For the following analysis, variables representing proportions where arcsin transformed, while the distance was log-transformed.

### 2.3. Statistical analysis

We first explored co-occurrence patterns among species across the study area using the probabilistic species co-occurrence models developed by Veech (2013). These models use the hypergeometric distribution to calculate the probability that two species spatially co-occur either less or more often than expected based on their mean incidence probabilities (Griffith, Veech, & Marsh, 2016). Using the observed co-occurrence frequencies and a specified alpha level (in this case, α = 0.05), species co-occurrences were classified as significantly positive or negative, or occurring at random.

Aside from the null model-based analyses, we explored through a multivariate analysis the potential association between the carnivore community composition and the variation in the site variables. First, we performed a non-metric multidimensional scaling (NMDS) on a similarity matrix constructed with Hellinger-transformed abundance data using the Bray-Curtis dissimilarity measure (Legendre & Legendre, 2012). The Hellinger transformation divides each cell by the total abundance at a site and then takes the square root (therefore reducing the effect of extremely abundant species), and has previously been shown to have desirable properties in the context of ordination (Legendre & Gallagher, 2001). Carnivore abundance was estimated through the relative abundance index (hereafter RAI; Carbone et al., 2001) by calculating the number of independent photographs divided by the total number of trap nights. This index was also used as an indicator of the intensity of habitat use by each species at each site. We consider as independent event those photographs taken with a delay greater than 45 min between each other. The assemblage environment relationships were examined by a *post hoc* fitting of environmental vectors onto the ordination space (Borcard, Gillet, & Legendre, 2018; Kindt & Coe, 2005). The result is a group of vectors with endpoint showing the direction of the environmental gradient, and a length that is proportional to the correlation between the environmental variable and the ordering of the NMDS. Beside the stress value, we checked the quality of the NMDS through Shepard plot and the significance of fitted vectors was assessed using a permutations test (n=999).

Additionally, and to take into account heterogeneous detection probabilities, we fit single-species single-season occupancy models to our data (MacKenzie et al., 2002, 2017). To avoid fitting a large number of models relative to the size of our dataset, we used the most influential variables resulted from the NMDS analysis as covariates into our occupancy models. Additionally, we used Hellinger-transformed RAI as covariable to evaluate potential effect of the capture rate of a given species on the probability of occupancy of other species. We carried out a model selection procedure and used the Akaike Information Criterion corrected for small sample sizes (AICc) to select supported models from sets of candidate models (Birochio, 2008; Burnham & Anderson, 2002). We considered being most supported those models with the lowest AICc score and the smallest number of parameters within 2 AICc units of the lowest AICc score. We conducted the following modeling procedure for each species: we first modeled the probability of detection by keeping the occupancy parameter constant and allowing detection to vary as a function of the covariates previously mentioned. For model selection, we considered all subsets and used AICc to identify the most supported model. We retained the most supported model to serve as the detection parameter for all subsequent models for that species. After that, to model the probability of occupancy, we built an initial additive global model consisting of the retained detection parameter and linear terms for each site-level covariate, we evaluated all subsets, and we considered the most supported model as our top model. To assess fit of each top model, we used a MacKenzie-Bailey goodness-of-fit test with parametric bootstrapping employing 1,000 simulations to approximate the distribution of the test statistic (MacKenzie & Bailey, 2004).

To study the daily activity patterns of carnivore species, we used the information on record time saved on each camera trap photograph to construct and visually evaluate circular plots representing the distribution of records along the 24-h cycle for each species and then tested for non-uniformity using the Rayleigh test (Jammalamadaka & SenGupta, 2001). Additionally, to evaluate the level of temporal overlap between species we fitted a Kerneldensity estimation that describes the temporal activity of each species and then calculated a coefficient of overlap in pairwise comparisons. As suggested by Ridout and Linkie (2009), we used the Δ1 estimator for those cases where the smallest number of photographic records was less than 50. This coefficient estimates the level of overlap between two activity distributions, and its value range from 0 for no overlap, to 1 for identical distributions. Given the small amount of records obtained for *G. cuja* and *C. chinga*, we only studied the temporal overlap between *L. geoffroyi* and *L. gymnocercus*. We obtained confidence intervals for each estimator using a bootstrap procedure (Linkie & Ridout, 2011).

All statistical analysis were performed in R (version 3.6.1) (R Development Core Team, 2013), using the package “cooccur” (Griffith et al., 2016), “unmarked” (Fiske & Chandler, 2011), “MuMIn” (Bartoń, 2013) and “overlap” (Meredith & Ridout, 2017).

## 3. Results

Total sampling effort was 1414 camera trap days and the mean effort per camera/sampling station was 48.75 days (range = 46-51 days). We obtained 83 independent records of carnivore species. The carnivore community of this area was formed by five species: the Pampas fox *Lycalopex gymnocercus*, Geoffroy’s cat *Leopardus geoffroyi*, Molina’s hog-nosed skunk *Conepatus chinga*, puma *Puma concolor*, and Lesser grison *Galictis cuja*.Although the jaguarundi *Puma yagouaroundi* and Pampas cat *Leopardus colocolo* are presumed to occur in this region, these felids were not detected in our survey. The following non-carnivore mammal species were also found in the “Arroyo Los Gauchos Nature Reserve”: the European red deer *Cervus elaphus*, European hare *Lepus europaeus*, wild boar *Sus scrofa* (three introduced species), big hairy armadillo *Chaetophractus villosus*, and several small rodents. It is also worth mentioning the presence of the Greater Rhea *Rhea americana*, a large bird that is listed as Vulnerable in Argentina (Navarro & Martella, 2011). *Lycalopex gymnocercus* and *L. geoffroyi* were the species with greatest RAI (3.68 and 1.49 independent events × 100 day^-1^, respectively), followed by *C. chinga* (0.28), *P. concolor* (0.21), and *G. cuja* (0.21) (Table 1).

**Table 1.** Summary of the number and percentage of independent photographic events, the capture rate (RAI) for the carnivore species found in a coastal area of Buenos Aires province (Argentina), and the number and percentage of sites were each one was detected.

| Species | N° photographic events (%) | RAI (N° photographic events × 100 day<sup>-1</sup>) | N° of sites with positive detection (%) |
| --- | --- | --- | --- |
| *L. gymnocercus* | 52 (62) | 3.68 | 17 (58.6) |
| *L. geoffroyi* | 21 (25) | 1.49 | 10 (34.5) |
| *C. chinga* | 4 (5) | 0.28 | 3 (10.3) |
| *P. concolor* | 3 (4) | 0.21 | 2 (6.9) |
| *G. cuja* | 3 (4) | 0.21 | 3 (10.3) |

*Leopardus geoffroyi* and *C. chinga* co-occurred at 3 sites, an amount significantly higher than the expected 1 station (Pgr = 0.033, Table 2). Similarly, *L. geoffroyi* and *L. gymnocercus* co-occurred at 8 sites while they were expected to co-occur at 5.9 sites; however, in this case, the p-value was only marginally significative (Pgr = 0.096, Table 2). The rest of the species paired combinations did not show any differences with a theoretical random pattern of co-occurrence.

**Table 2.** Pairwise comparison of the probability of spatial co-occurrence for all five carnivore species combinations in a coastal area of Buenos Aires province, Argentina. Values of P_lt_ < 0.05 indicate spatial segregation while values of P_gt_ < 0.05 indicate a positive association. Sites A and B refers to the number of camera-trap stations where Species A and B were detected, respectively.

| Sp. A | Sp. B | Observed co-occurrence | Probability of co-occurrence | Expected co-occurrence | $P_{lt}$ | $P_{gt}$ |
| --- | --- | --- | --- | --- | --- | --- |
| *L. geoffroyi* | *L. gymnocercus* | 8 | 0.202 | 5.900 | 0.984 | **0.096** |
| *L. geoffroyi* | *C. chinga* | 3 | 0.036 | 1.000 | 1.000 | **0.033** |
| *L. geoffroyi* | *P. concolor* | 1 | 0.024 | 0.700 | 0.889 | 0.579 |
| *L. geoffroyi* | *G. cuja* | 0 | 0.036 | 1.000 | 0.265 | 1.000 |
| *L. gymnocercus* | *C. chinga* | 3 | 0.061 | 1.800 | 1.000 | 0.186 |
| *L. gymnocercus* | *P. concolor* | 1 | 0.040 | 1.200 | 0.665 | 0.837 |
| *L. gymnocercus* | *G. cuja* | 3 | 0.061 | 1.800 | 1.000 | 0.186 |
| *C. chinga* | *P. concolor* | 0 | 0.007 | 0.200 | 0.800 | 1.000 |
| *C. chinga* | *G. cuja* | 0 | 0.011 | 0.300 | 0.712 | 1.000 |
| *P. concolor* | *G. cuja* | 0 | 0.007 | 0.200 | 0.800 | 1.000 |

**Table 3.** Occupancy models within 2 Akaike’s Information Criterion corrected for small sample size (AICc) units of top models, which show the strongest relationship between species occurrence and measured covariates in a coastal area of Buenos Aires, Argentina; p: detection parameter, ψ: occupancy parameter, K: number ofparameters in the model; MacKenzie-Bailey goodness-of-fit parameters are included for the top model of each species, and include X^2^, *P*-value, and *ĉ* as the overdispersion parameter. TG: tall grassland, LG: low grassland, BG: bare ground, J: *Juncus spp*., C: *Cortaderia spp.*, H: *Hyalis argentea*, D: distancie to shoreline, S: shrubs, RAI_LGy: relative abundance index of *Lycalopex gymnocercus*, RAI_LGe: relative abundance index of *Leopardus geoffroyi*.

| Species | Model | K | AICc | ΔAIC | Weight | $X^2$ | P-value | $\hat{c}$ |
| --- | --- | --- | --- | --- | --- | --- | --- | --- |
| *L. gymnocercus* | p(J+D) $\psi$(J+TG) | 6 | 174.8 | 0.00 | 0.206 | 281.59 | 0.143 | 1.36 |
| | p(J+D) $\psi$(RAI_Lge+TG) | 6 | 175.3 | 0.58 | 0.154 | | | |
| | p(J+D) $\psi$(.) | 4 | 176.2 | 1.45 | 0.100 | | | |
| | p(J+D) $\psi$(D+TG) | 6 | 176.2 | 1.47 | 0.099 | | | |
| *L. geoffroyi* | p(J) $\psi$(J+TG) | 5 | 112.6 | 0.00 | 0.260 | 366.70 | 0.051 | 2.46 |
| | p(J) $\psi$(J+TG+D+RAI_Lgy) | 7 | 112.8 | 0.15 | 0.241 | | | |
| | p(J) $\psi$(TG+D+RAI_Lgy) | 6 | 113 | 0.43 | 0.210 | | | |
| | p(J) $\psi$(J+D+RAI_Lgy) | 6 | 114 | 1.37 | 0.131 | | | |
| *C. chinga* | P(.) $\psi$(RAI_Lgy) | 3 | 38.2 | 0.00 | 0.428 | 51.58 | 0.267 | 0.89 |
| | P(.) $\psi$(D+RAI_Lgy) | 4 | 39.7 | 1.47 | 0.205 | | | |
| *G. cuja* | P(.) $\psi$(J) | 3 | 33.3 | 0.00 | 0.376 | 6.90 | 0.822 | 0.17 |
| | P(.) $\psi$(J+RAI_Lgy) | 4 | 34.6 | 1.35 | 0.192 | | | |
| | P(.) $\psi$(J+D) | 4 | 35.2 | 1.97 | 0.140 | | | |
| *P. concolor* | P(D) $\psi$(D) | 4 | 26.3 | 0.00 | 0.693 | 54.60 | 0.730 | 0.536 |

The NMDS analysis showed a stress value of 0.028 and the Shepard plot did not show discrepancies between the original dissimilarities and the multidimentional scaling solution (R^2^ = 0.99). Axes were significantly related to the distance to shoreline (p < 0.05) and marginally so to the proportion of tall grassland (p = 0.096) (Figure 2). *Leopardus geoffroyi* appeared to be the species more strongly associated to sites near the shoreline and with higher proportion of tall grassland and, to a lesser extent, *Cortaderia spp*.; whereas *L. gymnocercus* tended to use the most inland sites, relatively farther from the shoreline and with higher proportion of *H. argentea* and low grassland, and a smaller proportion of tall grassland. *Puma concolor* appeared to be strongly associated to sites with higher proportion of *Juncus spp*. and tall grassland, a smaller proportion of *Cortaderia spp.*, and relatively far from the coast. *Conepatus chinga* habitat use was related to sites with small proportion of *Juncus spp*. and high percentage of *Cortaderia spp*. and, to a lesser extent, bare ground. Finally, *G. cuja* showed an association to sites not close to the shoreline, with higher proportion of *Juncus spp*. and tall grassland, and less *Cortaderia spp*.

**Figure 2.**
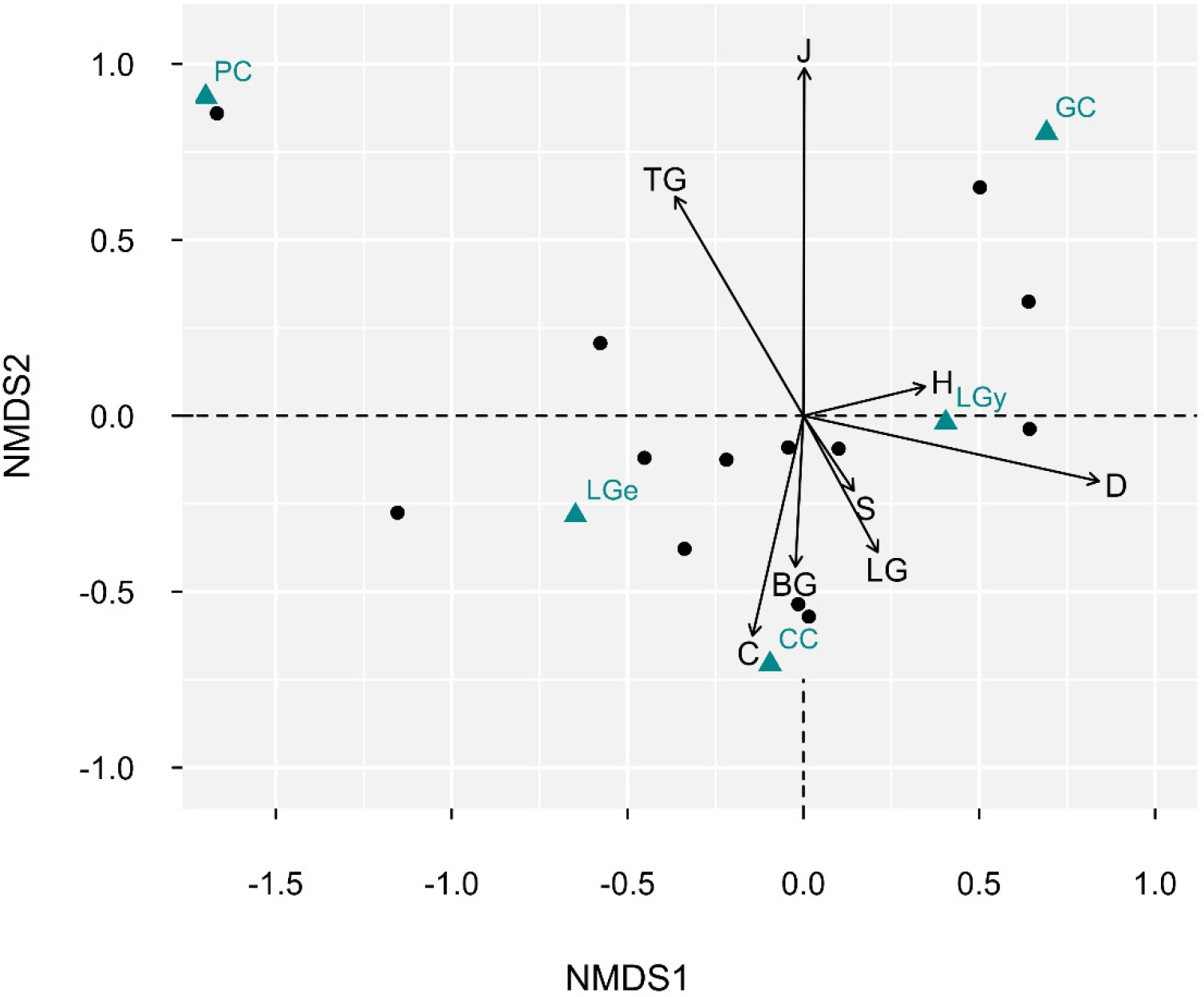
Result of Non-Metric Multi-dimensional Scaling for the carnivore species found in a coastal area of Buenos Aires province (Argentina) and the *post hoc* environmental vectors fitting to ordination (Stress = 0.028). Vectors show the direction of linear correlation of the environmental variables with ordination scores. TG: tall grassland, LG: low grassland, BG: bare ground, J: *Juncus spp*., C: *Cortaderia spp.*, H: *Hyalis argentea*, D: distancie to shoreline, S: shrubs, LGe: *Leopardus geoffroyi*, LGy: *Lycalopex gymnocercus*, CC: *Conepatus chinga*, PC: *Puma concolor*, GC: *Galictis cuja*.

**Figure 3.**
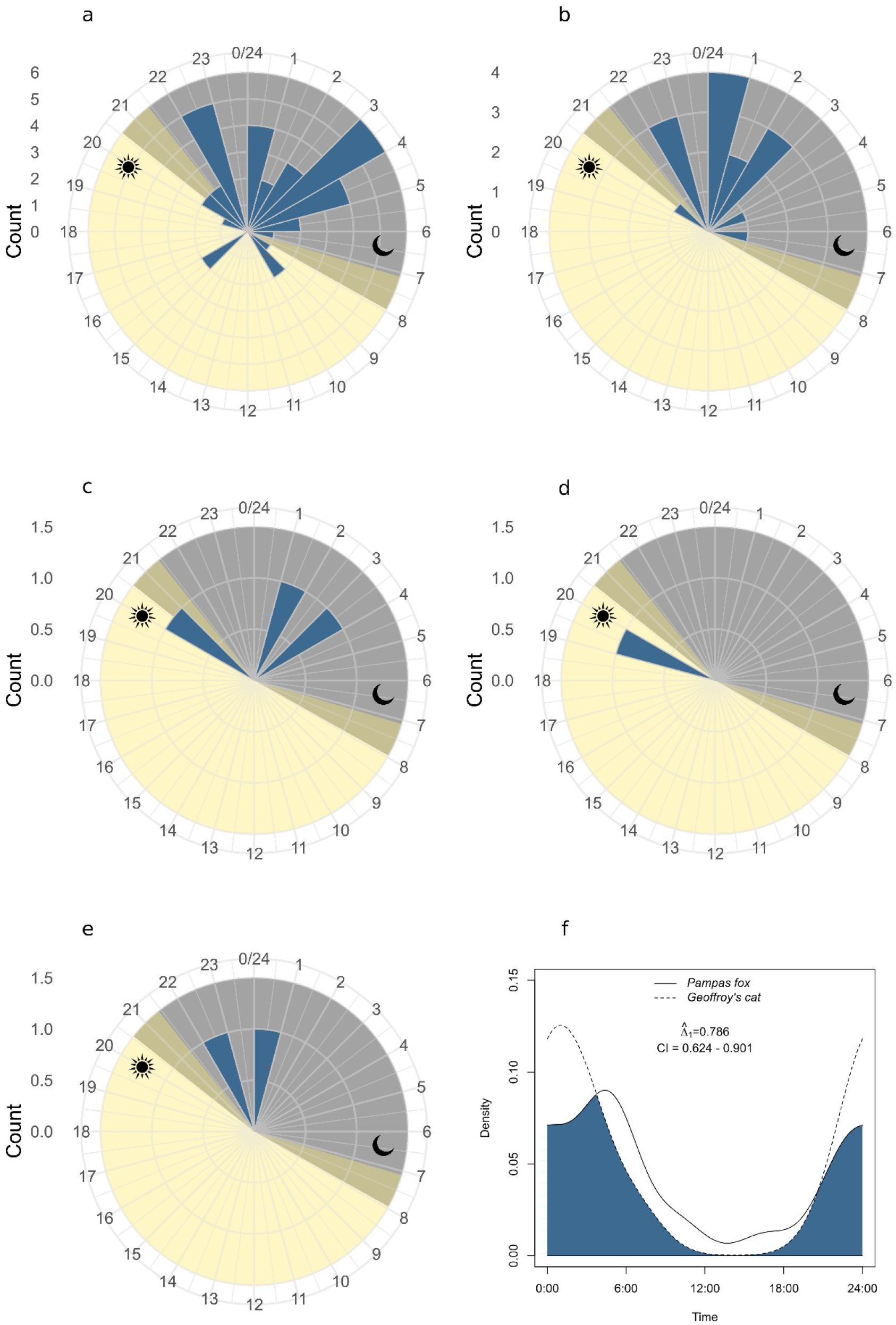
Activity patterns of carnivore species in a coastal area of Buenos Aires province, Argentina: a) *Lycalopex gymnocercus*, b) *Leopardus geoffroyi*, c) *Conepatus chinga*, d)*Galictis cuja*, e) *Puma concolor*; and f) pairwise comparison of daily activity patterns between *Lycalopex gymnocercus* and *Leopardus geoffroyi*. Δ_1_: overlap coefficient; CI: confidence interval.

The top occupancy models for *L. gymnocercus, C. chinga, G. cuja*, and *P. concolor* exhibited evidence of model fit (P > 0.05), whereas lack of fit (p ≈ 0.05) and overdispersion (*ĉ* > 2) was found in the top model for *L. geoffroyi* (Table 4). The top models for both Pampas fox and Geoffroy’s cat included a negative logistic relationship with the proportions of *Juncus sp*. and of tall grassland for the occupancy parameter (Table 4). The detection probability for Pampas foxes was negatively related to the proportion of *Juncus sp*. and positively to the distance to the shoreline, while it was positively related to *Juncus sp*. for Geoffroy’s cat. A negative effect of the distance from the shoreline on the probability of occupancy of this felid was found in three of the four models included in the set with ΔAICc < 2. Although it was not the case for the top model, the occupancy probability of Geoffroy’s cat showed a positive relationship with the occupancy probability of Pampas fox, and vice versa. Top models for both *C. chinga* and *G. cuja* did not support any variable for the detection parameter, but showed a logistic positive relationship with the capture rate of Pampas fox (for the case of *C. chinga*) and a positive relationship with the proportion of *Juncus sp*. (for the case of *G. cuja*). *Puma concolor* was the only species with a single model in the set of models with AICc <2 that showed a negative relationship with distance to the shoreline for the detection parameter and a positive relationship with the same variable for the occupancy parameter.

Carnivores were mostly nocturnal, except for *G. cuja*, which was only captured during daytime (Figure 4). All records of the *P. concolor* and almost all those of *C. chinga* were obtained at night. The Pampas fox seemed to have a bimodal pattern with activity concentrated between 22:00 – 23:00 and 3:00 – 4:00. The highest rate of increase in activity was at dusk (approximately at 21:00) and only few pictures were recorded during daytime (21.6% of the independent events). Most of the events of the Geoffroy’s cat were obtained during nighttime, with a concentration around midnight (0:00 – 1:00). There were no records during daytime and a few in crepuscular periods. Pampas fox and Geoffroy’s cat showed a relatively similar activity patterns (Δ1 = 0.786; CI = 0.624 – 0.901).

## 4. Discussion

With a relatively limited surveying effort we were able to detect five of the seven species of Mammalian carnivores that may potentially occur in the coastal area of the southern part of Buenos Aires province. Although we were not able to record the presence of the Pampas cat and the jaguaroundi, we cannot exclude the possibility that they simply went undetected. Based on the presence of Pampas cats in the sandy dune areas found less than 200 km away in the Espinal ecoregion of Buenos Aires province (Caruso, Manfredi, Vidal, Casanave, & Lucherini, 2012) and other records in proximity of the city of Bahia Blanca (located aprox. 100 km away), this possibility appears more likely for this felid than the jaguaroundi. There are no recent records of the latter in the region and its population densities in the southernmost part of Buenos Aires province appear to be very low (Luengos Vidal, Guerisoli, Caruso, & Lucherini, 2017). A comparison with the previous information available suggests that in neighboring Pampas areas, which have been more intensively affected by anthropogenic modifications, some of the species we recorded are rare, such as the Geoffroy’s cat (Castillo et al., 2008), or in a process of recolonization, such as the puma (Chimento & De Lucca, 2014). Furthermore, the capture rates (RAI) found in our study area indicates that Geoffroy’s cats are similarly abundant and pumas are even more common compared to a protected area (Reserva Natural de Objetivo Definido Laguna de Chasicó) with relatively preserved woodland typical of the Espinal ecoregion of Buenos Aires province (Benzaquín, 2008; Caruso et al., 2012).

Both the RAI values and the naïve occupancy (i.e., the proportion of sampling site where a species is recorded) showed that the Pampas fox is the most common carnivore species in the coastal area of the southern part of Buenos Aires province, followed by the Geoffroy’s cat. The numeric dominance of Pampas fox is not surprising, because this canid is a very adaptable carnivores, capable of opportunistically feed on a variety of prey (e.g., Bossi, Migliorini, Santos, & Kasper, 2019; Castillo, Birochio, Lucherini, & Casanave, 2011; Farias & Kittlein, 2008) and still relatively common in the agroecosystems of the Pampas. Conversely, the relatively high frequency of records and occupancy of the Geoffroy’s cat in our study area, especially when compared to those of *C. chinga* and *G. cuja*, was unexpected, based on the available information on the present-day Pampas landscapes. Although caution is required when comparing data that do not account for differences in detection probability, the information recorded previously in the Argentine Pampas indicate that *C. chinga* is the second most abundant carnivore both in natural mountain grasslands and croplands (Luengos Vidal, 2009). *C. chinga* was also found to be very abundant in the Brazilian Pampas, where this Mephitid could even become the most common carnivore in intensive agriculture areas and ranchlands, while the Geoffroy’s cat was much rarer (Kasper et al., 2012). Thus, our findings suggest that the “Arroyo Los Gauchos Nature Reserve” can be conserving habitats where the carnivore community is able to maintain relatively high densities of species that are comparatively more sensitive to human interference such as the Geoffroy’s cat and the puma.

Because Pampas foxes readily eats carrion and have been reported to prey on several animals associated to saline water, such as crabs, crustaceans, and even fish (García and Kittlein, 2005; Lucherini and Luengos Vidal, 2008; Bossi et al., 2019), these canids could be expected to use the areas close to the shoreline more intensively than other carnivores. We found little support for this hypothesis. Only one of the top four models included a negative relationship between the distance from the shoreline and the Pampas fox occupancy. Additionally, the NMDS analysis showed that *L. gymnocercus* tended to use more intensively sites at greater distances from the ocean. However, the finding that the probability of detecting this canid also increases with increasing distance from the shoreline may indicate that the NMDS results are affected by this greater detection. Also contrary to our expectations, the negative effect of the distance from the shoreline on the probability of occupation appeared to be stronger for the Geoffroy’s cat, which is in accordance also with the results of the NMDS analysis. There are several possible explanations for these findings, but we argue that the spatial distribution of resource availability is a mayor candidate and a factor that should be investigated for a better understanding of the dynamics carnivore habitat use in this ecosystem.

In agreement with our initial hypothesis - and congruently with the results of the occupancy analysis -, we found evidence of a lack of spatial and, to a lower extent, temporal avoidance between the two most common carnivore species of our study area. These results are similar to those found by a previous study that specifically addressed the mechanisms of coexistence between *L. geoffroyi* and the culpeo *Lycalopex culpaeus*, a fox slightly larger than the Pampas fox (Gantchoff & Belant, 2016). These authors found high spatio-temporal overlap between these two carnivores and concluded that diet segregation was the most likely mechanism favoring their coexistence in the Andean forest of Patagonia (Gantchoff & Belant, 2016). Based on the available information on the trophic niche of the Geoffroy’s cat and the Pampas fox, in general and in areas where they co-occur (Kasper, Peters, Christoff, & de Freitas, 2016), we argue that dietary segregation, coupled and possibly also made possible, by the diversity and abundance of food resources found in our study site, may be the major factor facilitating the coexistence of these two carnivores in the coastal areas of Buenos Aires province.

## 5. Conclusions

Our findings support those from Bilenca y Miñarro (Bilenca & Miñarro, 2004) and (Monserrat & Celsi, 2009; Monserrat et al., 2012) indicating that the coastal dunes have an important role in the conservation of the biodiversity of Buenos Aires province. They protect not only endemic species with very limited distributions but also offer refuge to species that were probably more common in the natural ecosystems typical of the Pampas grassland before they were heavily transformed by anthropogenic activities. This conclusion is supported by the relatively diversified and abundant guild of vertebrates, especially carnivores, that we found. It is clear that the vertebrate community of the coastal areas of Buenos Aires cannot be considered pristine, not only because they are typically used as grazing areas for extensive livestock activities but also because of the presence of several introduced species. However, livestock has not made irreversible damages yet and the presence of exotic prey may reduce the potential conflicts between carnivores (namely, *P. concolor* and *L. gymnocercus*) and ranchers over predation on livestock, which are intense in Buenos Aires province (De Lucca, 2010; Guerisoli et al., 2017). Although more information is certainly required, we conclude that the coastal areas of the southernmost portion of Buenos Aires deserve protection. This does not necessarily imply a complete halt to tourism development, the activity that currently seems the strongest and most promising economic activity for these coasts. Wildlife conservation is compatible with carefully-designed ecotourism and limited infrastructure development and this may be a unique chance for the areas of Buenos Aires coast that have not been affected yet by poorly planned, conservation-unfriendly urbanization.

## Acknowledgment

This work was financially supported by Secretaría General de Ciencia y Técnica - Universidad Nacional del Sur (PGI 24/B152, 24/B198 and 24/B243) to EBC and Université de Sherbrooke (Maîtrise en Biologie - Écologie Internationale). M.C.M. and M.S.A. were founded by a postdoctoral and a doctoral scholarship, respectively, from the Consejo Nacional de Investigaciones Científicas y Técnicas (CONICET). We would like to express our special gratitude to the ranch owners surrounding the study area who kindly provided us with logistics support in the field. We also thank to V. Robichaud, P. Costilla and G. Zapperi who collaborated with fieldwork, data entry and statistical analysis.

